# Chromatin accessibility of primary human cancers ties regional mutational processes with tissues of origin

**DOI:** 10.1101/2021.05.14.444202

**Authors:** Oliver Ocsenas, Jüri Reimand

**Affiliations:** Computational Biology Program, Ontario Institute for Cancer Research, Toronto, ON, Canada; Department of Medical Biophysics, University of Toronto, Toronto, ON, Canada; Department of Molecular Genetics, University of Toronto, Toronto, ON, Canada

## Abstract

**Background:** Regional distribution of somatic mutations in cancer genomes associates with DNA replication timing (RT) and chromatin accessibility (CA), however normal tissues and cell lines have contributed these insights while associations with the epigenomes of primary cancers remain uncharacterized.

**Results:** Here we model megabase-scale mutation burden in whole cancer genomes using ∼900 CA and RT profiles of primary cancers, normal tissues, and cell lines. CA profiles of primary cancers, rather than normal tissues, predict regional mutagenesis in most cancer types. Regional mutation burden associates with the CA profiles of matching cancer types, indicating tissue-specific determinants of mutagenesis. However, mutagenesis in squamous cell and lymphoid cancers instead associates with RT profiles. Mutational signatures also show tissue-specific associations with cancer epigenomes, especially for carcinogen-induced and unannotated signatures. Lastly, while each cancer type includes certain frequently-mutated genomic regions exceeding epigenome-informed predictions of mutation burden, these regions show a pan-cancer convergence to biological processes involved in development and cancer. Thus, modelling excess mutations using epigenomes highlights known cancer driver genes as well as frequently mutated non-coding regions.

**Conclusions:** The dominant association of regional mutation burden with cancer epigenomes suggests that many passenger mutations are determined by the epigenetic landscapes of transformed cells and may occur later in tumor evolution. CA-informed models help find cancer genes and pathways with positive selection and highlight regions where additional mutation burden is contributed by local mutational processes. This study underlines the complex interplay of mutational processes, genome function and evolution in cancer and tissues of origin.

## INTRODUCTION

The cancer genome is a footprint of its evolution and molecular environment that is shaped by somatic mutations such as single nucleotide variants (SNVs) and structural alterations ^1,2^. Cancer initiation and progression is caused by a small number of driver mutations that provide cells with selective advantages ^3-5^, however most mutations are functionally neutral passengers that are caused by various mutational processes ^6-8^. Somatic mutations also occur in normal tissues and are frequently observed in known cancer genes ^9,10^. Thus, we need to understand mutational processes to decipher tumor etiology and evolution and better characterize driver mutations.

Mutational processes act at different scales of the cancer genome ^11,12^. Single base substitution (SBS) signatures affect certain trinucleotide context of DNA and are associated with aging, carcinogen exposures, defects in DNA repair pathways, and cancer therapies ^6,13^. At a 100-nucleotide resolution, local mutational processes disproportionately affect certain non-coding genomic elements such as transcription start sites and binding sites of gene-regulatory proteins such as CTCF ^14-16^. At the regional, megabase-scale resolution of the genome, mutation burden correlates with DNA replication timing (RT), chromatin accessibility (CA) and transcriptional activity ^17-19^. Early-replicating, transcriptionally active regions of open chromatin have fewer mutations than late-replicating, passive regions of heterochromatin, potentially due to increased error rates and decreased mismatch repair later in DNA replication ^20-23^. SBS signatures are also distributed asymmetrically with respect to DNA replication origins and timing ^24^. Regional mutation burden is associated with epigenetic information of related normal cells, providing evidence of cells of cancer origin contributing to somatic variation ^25^ and allowing classification of cancers of unknown origin ^26^. However, the precise molecular mechanisms driving these mutational processes remain incompletely understood. In particular, cell lines and normal tissues have been used to associate chromatin accessibility and mutation burden in cancer, while the epigenetic landscapes of primary human cancers remain unexplored.

Here we studied cancer epigenomes as determinants of regional mutagenesis in thousands of whole cancer genomes through a diverse collection of CA and RT profiles of cancers, normal tissues, and cell lines. CA profiles of matching cancer types, rather those than normal tissues, are the major determinants of regional mutagenesis and mutational signatures in most cancer types.

We found tissue-of-origin effects of CA and RT in most predictions, bespoke deviations in specific cancer types and mutational signatures, and a pan-cancer convergence of excess mutations to cancer driver genes and developmental pathways. Together, these results underline the spatial complexity of regional mutagenesis in cancer genomes and highlight epigenome-informed avenues to discover driver mutations.

## RESULTS

### Chromatin accessibility of primary cancers is a major determinant of regional mutagenesis

To study cancer epigenetic profiles as determinants of regional mutagenesis in cancer genomes, we collected 773 ATAC-seq profiles of genome-wide CA measurements in primary human cancers, normal tissues and cell lines from ENCODE3, TCGA, and additional studies ^27-34^, as well as 96 RepliSeq profiles of DNA replication timing measurements in 16 cell lines and six cell cycle phases ^35^ (**Figure 1A; Supplementary Figure 1, Supplementary Table 1**). As regional mutation burden, we studied 23 million SNVs in 2,465 highly-mappable genomic regions of one megabase mapped across 2517 whole cancer genomes of 37 cancer types of the ICGC/TCGA PCAWG project ^1^ (**Figure 1B**). The 869 CA and RT profiles were derived as mean signal intensity values per megabase.

**Figure 1.**
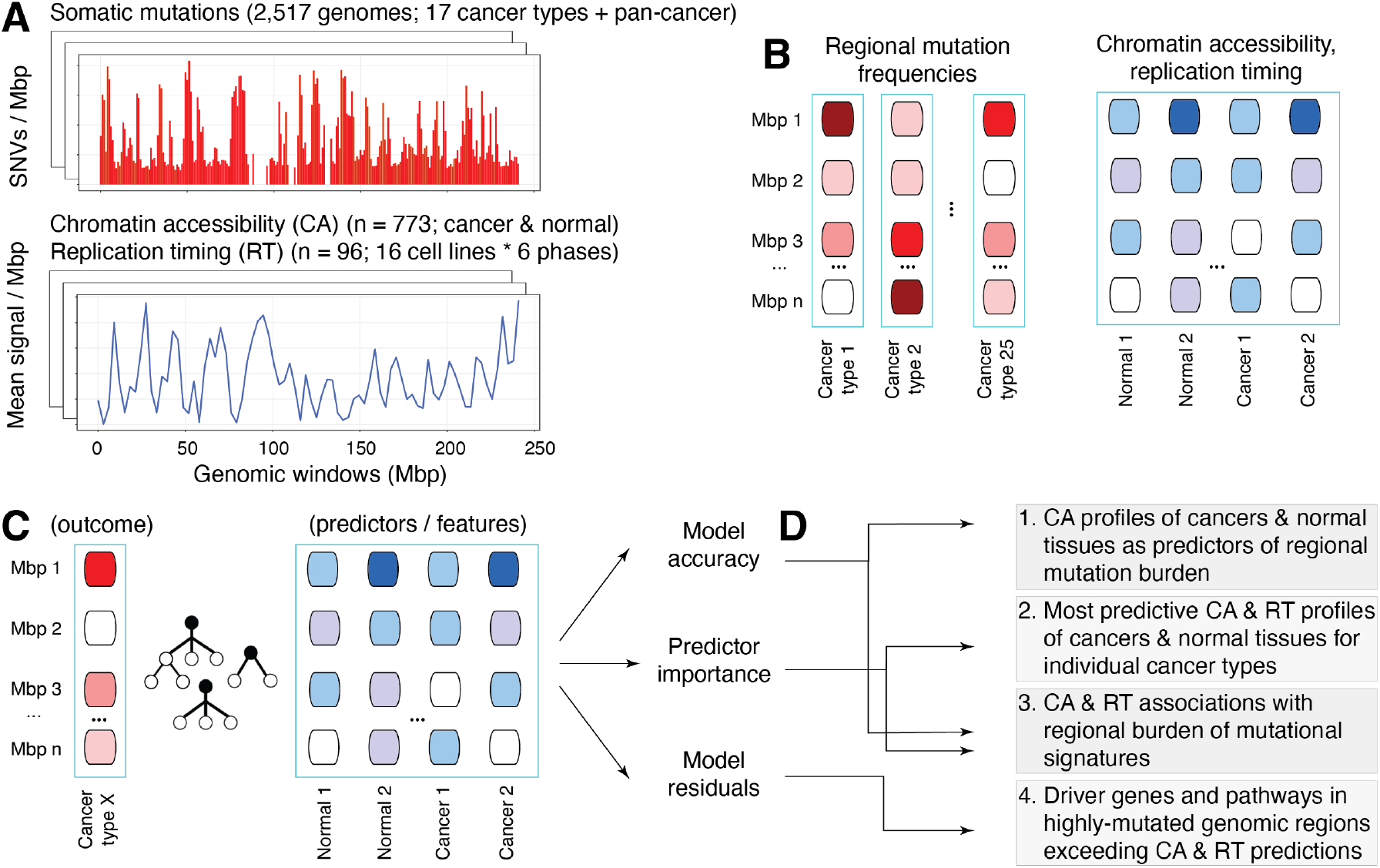
Characterizing chromatin accessibility (CA) and replication timing (RT) as determinants of regional mutagenesis in cancer genomes. **A**. Somatic mutations in cancer genomes (top) and CA and RT datasets of normal tissues and cancers (bottom) were integrated to study regional mutational processes. Somatic single nucleotide variants (SNVs) of 2,517 whole cancer genomes were analyzed with 869 genome-wide profiles, including 773 CA profiles of primary human cancers, normal tissues, and cell lines from ATAC-seq experiments, and 96 RT profiles of and six cell cycle phases in 16 cell lines from RepliSeq experiments. **B**. Genomic regions of one megabase (Mbp) were analyzed. Regional mutation burden was estimated as the number of SNVs per megabase region. The mean values scores per region were derived for CA and RT profiles. **C**. Random forest models were trained using regional mutation burden profiles as the outcome and CA and RT profiles as the predictors. We analyzed the pan-cancer dataset and 17 datasets of specific cancer types with relevant CA and RT profiles available. **D**. To associate regional mutation burden with CA and RT, mutational signatures, and cancer driver genes, the random forest models were evaluated in terms of accuracy, predictor importance, and model residuals.

To map the complex non-linear associations of CA and RT profiles with regional mutagenesis, random forest regression models were trained with mutation burden profiles as outcomes and CA and RT profiles as predictors (*i*.*e*., features) for the 17 cancer types for which both genomic and relevant epigenomic profiles were available (**Figure 1C**). The most informative predictors were quantified using statistical analysis and local feature prioritization of random forest models ^36^ (**Figure 1D**). As expected, genome-wide profiles of regional mutation burden clustered according to cancer types (**Supplementary Figure 2**).

We asked whether the CA profiles of cancers or those of normal cells and tissues showed stronger associations with regional mutational processes in cancer genomes. We predicted regional mutation burden in pairs of random forest models with matched data splits where the predictors included either CA profiles of primary cancers or CA profiles of normal cells and tissues, respectively. RT profiles were also included as predictors in both models to estimate the relative contributions of CA.

In most cancer types, CA profiles of primary cancers showed stronger associations with regional mutagenesis than CA profiles of normal cells and tissues (13 of 17, *P* < 0.05) (**Figure 2A**). The most pronounced signal was observed in breast cancer where the regional mutagenesis predictions informed by cancer CA profiles were nearly twice as accurate as those informed by CA profiles of normal tissues (median adj.R^2^ 0.70 *vs*. 0.38; *P* < 0.001) (**Figure 2B**). Stronger associations of cancer CA profiles and regional mutagenesis were also found in cancers of the prostate, uterus, and kidney, and melanoma: the improvement in prediction accuracy was above 10% in those cancer types (*P* < 0.001). Stronger associations with cancer epigenomes were also confirmed in the pan-cancer analysis across 37 cancer types, with a small but statistically significant improvement in model accuracy (adj.R^2^ 0.90 *vs*. 0.88; *P* < 0.001). As the only exception, regional mutation burden in liver cancer better associated with CA profiles of normal tissues (adj.R^2^ 0.85 *vs*. 0.83; *P* = 0.044). The analysis provided inconclusive evidence for four cancer types including lymphoid cancers (BNHL, CLL) and lung and thyroid adenocarcinomas. We confirmed that the accuracy of regional mutation burden predictions in individual cancer types was not significantly correlated with the overall mutation burden or the number of sequenced genomes per cancer cohort (**Supplementary Figure 3**).

**Figure 2.**
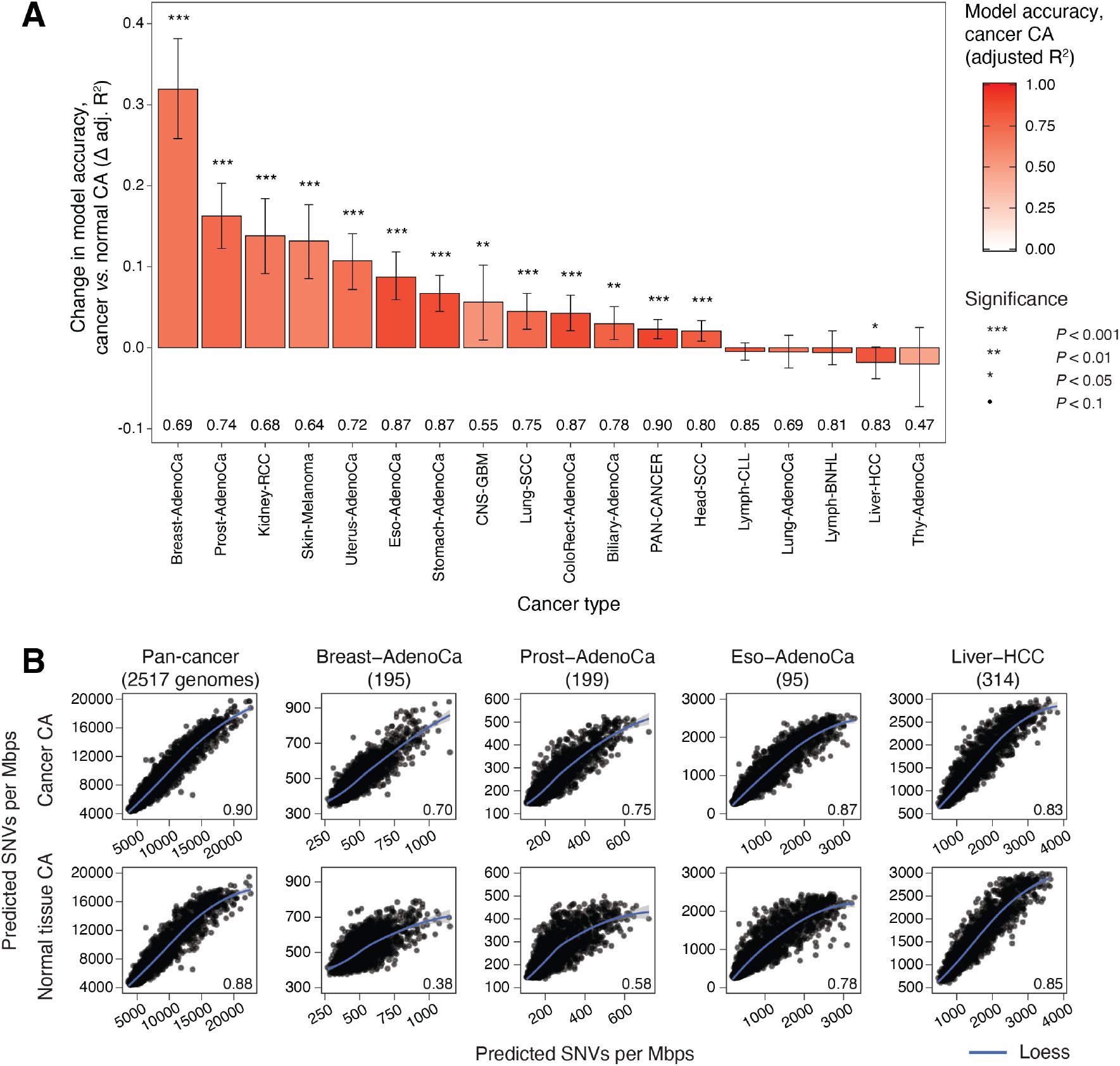
Chromatin accessibility of primary cancers is a major determinant of regional mutagenesis in cancer genomes. **A**. Random forest models informed by CA profiles of primary cancers are more accurate predictors of regional mutation burden, compared to models informed by CA of normal tissues. Bar plot shows relative change in prediction accuracy (Δ adjusted R^2^) of random model regression models informed by CA profiles of primary cancers, compared to matching models informed by CA of normal tissues. Replication timing (RT) profiles are included in all models as reference. Empirical P-values and 95% confidence intervals from bootstrap analysis are shown. Accuracy values of models informed by cancer CA profiles are listed below the bars (adjusted R^2^). **B**. Examples of regional mutation burden predicted using models informed by CA profiles of cancer (top) *vs*. CA profiles of normal tissues (bottom). Scatterplots show model-predicted and observed mutation burden (X *vs*. Y-axis) in one-megabase regions. Prediction accuracy values are shown (bottom right).

In summary, this analysis shows that in most cancer types, regional mutagenesis is more strongly associated with chromatin accessibility of primary human cancers rather than normal tissues and cell lines, even when accounting for DNA replication timing in the comparison. The diverse collection of epigenomes included as predictors suggests that tissue-specific chromatin features of individual cancer types, as well as pan-cancer chromatin features of proliferative cells may contribute to regional mutagenesis.

### Top predictors of regional mutagenesis match cancer types and sites of origin

To interpret the determinants of regional mutagenesis, we asked which specific CA and RT profiles contributed the most to the predictive models when using all 869 cancer and normal epigenomes as predictors. We selected the five most significant predictors for each cancer type (permutation *P* < 0.001). These 85 CA and RT profiles were quantified using Shapley additive explanation (SHAP) scores ^36^ that measured the directional associations of individual profiles with the regional mutation burden in all genomic regions. As expected, regional mutation burden negatively correlated with CA profiles of primary cancers and normal tissues (ρ_cancer_ = -0.74 vs ρ_normal_ = -0.79; *P* < 10^−16^) (**Figure 3A**). A dual relationship was apparent in RT profiles: RT profiles of late cell cycle phases positively correlated with regional mutation burden while RT profiles of early cell cycle phases correlated negatively (ρ_late_ = 0.75 *vs*. ρ_early_ = -0.84, *P* < 10^−16^). The inverse relationships of CA and RT with respect to regional mutation burden are consistent with previous studies ^17-23^ and extend here to a diverse collection of epigenomes from primary cancers and normal tissues.

**Figure 3.**
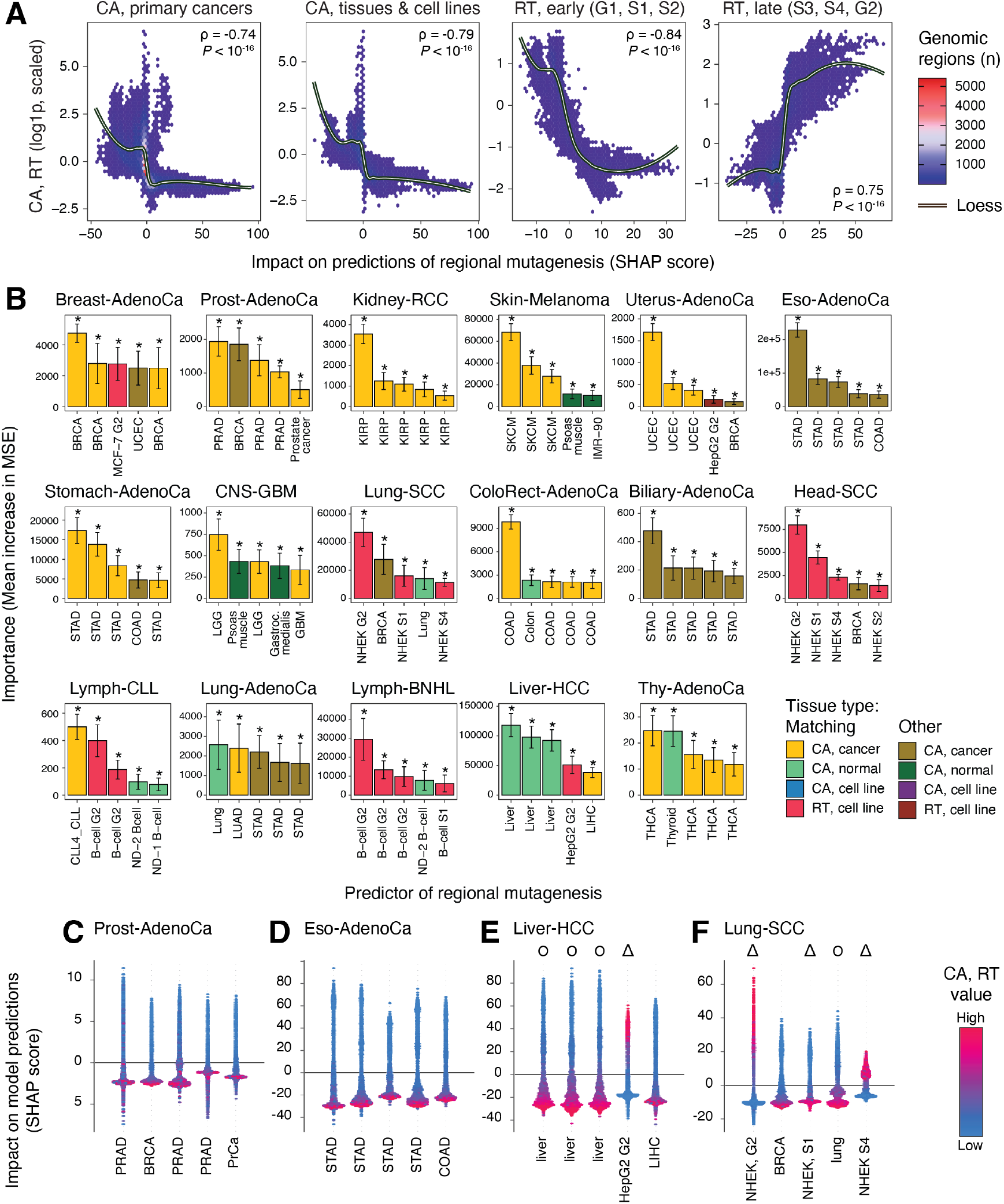
Top predictors of regional mutagenesis tie cancer types and sites of origin. **A**. Quantitative genome-wide associations of regional mutation burden with the most informative CA and RT profiles in random forest models. 2D-density plots show the association CA and RT scores (Y-axis) and Shapley feature importance (SHAP) scores in each genomic window (X-axis) across all cancer types. CA profiles for cancer and normal samples, early RT profiles, and late RT profiles are plotted separately. CA and early RT profiles negatively correlate with regional mutation burden while late RT profiles correlate positively. Spearman correlation values are shown (top right). **B**. CA profiles of primary cancers are the top predictors of regional mutagenesis in most cancer types. Bar plot shows the importance scores of the five most important predictors of random forest models for 17 cancer types (permutation *P* < 0.001). Error bars show ±1 standard deviation from bootstrap analysis. Brighter colors indicate the predictors where the epigenomic profile (CA or RT) matches the mutation profile of the related cancer type. **C-F**. Top predictors of regional mutation burden in individual cancer types. Shapley additive explanation (SHAP) scores show the impact of a given predictor on the predictions of regional mutation burden (Y-axis) relative to the values of the predictor (CA or RT; color gradient). In CA profiles, higher values (red) primarily associate with lower SHAP scores corresponding to increased mutation rates, while in contrast, higher values in late RT profiles associate with higher SHAP scores. Symbols indicate RT profiles (triangles) and CA profiles of normal tissues (circles).

We examined the most significant predictors of regional mutation burden for each cancer type (**Figure 3B**). CA profiles of primary cancers dominated among the strongest predictors of regional mutation burden in 12 of 17 cancer types. Most CA profiles represented the same or related cancer type where the regional mutation burden was predicted, underlying tissue-specific interactions of chromatin state and somatic mutagenesis Overall, CA profiles of primary cancers were enriched among the top predictors (55 of 85 profiles observed *vs*. 41 expected, Fisher’s exact *P* = 0.011), confirming the stronger association with primary cancer epigenomes and regional mutagenesis. Regional mutation burden measured in the breast, prostate, kidney, stomach, and thyroid cancer cohorts of the PCAWG WGS dataset associated with the CA profiles of the matching cancer samples in TCGA (BRCA, PRAD, KIRP, STAD, and THCA, respectively). For example, in prostate cancer, four CA profiles of primary prostate cancers and one breast cancer profile associated negatively with regional mutation burden (**Figure 3C**). Additional associations were apparent at the level of organ systems. CA profiles of stomach and colorectal cancers were the top predictors of regional mutation burden in biliary and esophageal cancers (**Figure 3D**), suggesting epigenetic and mutational similarities of cancers of the gastrointestinal tract. As another example, regional mutagenesis in in breast cancer genomes was significantly associated with one CA profile of uterine cancer, and a similar association with breast cancer CA was apparent in uterine cancer genomes, perhaps explained by common mutational processes in cancers of the female reproductive system. Therefore, regional mutational processes in individual cancer types have strong tissue-specific interactions with the epigenomes of these cancer types.

Fewer CA profiles of normal tissues were found among top predictors of regional mutation burden. The strongest association with normal tissue epigenomes was apparent in liver cancer, as three CA profiles of normal liver were detected as the highest-ranking predictors of regional mutation burden (**Figure 3E**). CA profiles of normal tissues were identified as predictors in seven other cancer types, however their feature importance scores were lower compared to CA profiles of related primary cancers. As expected, these CA profiles of normal tissues also matched the cancer types where regional mutagenesis was measured. For example, in thyroid cancer, one normal thyroid CA profile and four primary cancer CA profiles of the matching cancer type (THCA) were the top predictors of regional mutation burden.

Replication timing showed the strongest associations in squamous cell cancers (SCC) and lymphoid cancers, reflecting tissue-specific effects. Mutations in Lung-SCC and Head-SCC cohorts of PCAWG associated with RT profiles of NHEK cells, a squamous cell line of human epidermal keratinocytes (**Figure 3F**). Similarly, regional mutation burden in lymphoid cancers (Lymph-BNHL, Lymph-CLL) strongly associated with RT profiles of B-cells. One potential explanation of these normal cell lines associating with regional mutagenesis in those cancer types, rather than CA profiles of primary cancers, is an earlier occurrence of mutagenesis in the evolution of these cancer types. In the genomes of Lung-SCC and Head-SCC cohorts of PCAWG, many mutations are associated with signatures of tobacco exposure, while somatic hypermutation contributes to genome variation in normal B-cells and lymphomas ^37^.

Most RT predictors of regional mutagenesis we found in the analysis (12/15) represented late-replicating cell cycle phases G2 and S4. Individual RT profiles positively associated with regional mutation burden in late-replicating regions (*e*.*g*., phase G2 of MCF-7 in breast cancer) and negatively in early-replicating regions (*e*.*g*., phase S1 of HNEK in Head-SCC), consistent our analysis above (**Figure 3A**) and with earlier observations that elevated regional mutagenesis is caused by increased DNA damage and decreased repair in late-replicating regions ^20^.

Fewer RT profiles occurred among top predictors compared to CA profiles in other cancer types. RT profiles of matching cell lines (MCF-7, HepG2) were found among predictors of regional mutation burden in breast and liver cancer, respectively, and the latter RT profile was also a minor but significant predictor in uterine cancer. In general, fewer and less-diverse RT profiles of cell lines were available for this analysis, and these offer only a limited representation of mutational processes in different cancer types. In contrast, the larger set of CA profiles represents more cancer types and provides complementary information to RT. This analysis extends our findings of tissue-specific CA and RT profiles as the principal predictors of regional mutagenesis and underlines the effects of cell-of-origin and tumor heterogeneity. Dominance of cancer CA profiles among top predictors in most cancer types is consistent with our first observations that CA profiles of primary cancers provide accurate predictions of regional mutagenesis.

### Associations of mutational signatures with chromatin accessibility and replication timing

We asked whether the associations of regional mutagenesis with CA and RT are further explained by mutational signatures. We quantified the megabase-scale mutation burden separately for the major single base substitution (SBS) signatures based on PCAWG datasets ^6^ and predicted their regional distributions using random forest regression. We then selected the top five CA and RT profiles that most significantly associated with the burden of each mutational signature and cancer type (*P* < 0.001).

We compared the top CA and RT profiles that associated with total regional mutation burden and the burden of individual mutational signatures (**Figure 4A**). Top predictors of individual signatures were often consistent with predictors of bulk regional mutation burden and showed tissue-specific associations of mutagenesis and chromatin accessibility. Matching CA profiles of cancers were the top predictors of mutational signatures in breast, kidney, colorectal and stomach cancers, while RT profiles of matching cell lines associated with mutations in SCCs and lymphoid cancers. Top predictors of endogenous and exogeneous signatures were also mostly consistent, indicating that various mutational processes are affected by the epigenetic landscapes of cancers or their normal cells of origin.

**Figure 4.**
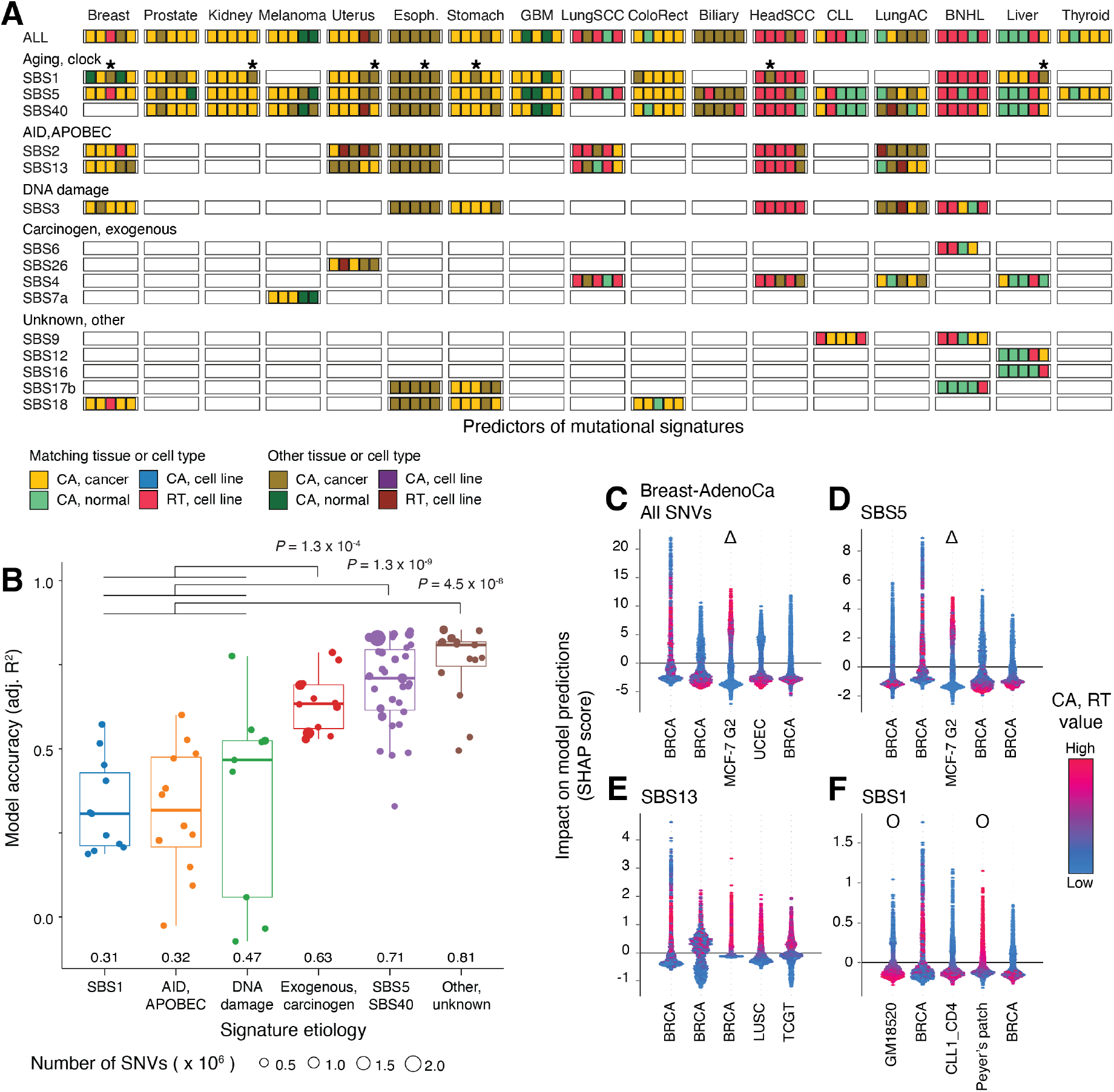
Associations of mutational signatures with chromatin accessibility and replication timing. **A**. Top predictors of megabase-scale mutation burden of single base substitution (SBS) signatures (top five predictors; *P* < 0.001, permutation test). Colors indicate the predictor type (CA, RT) and its relationship to the cancer type where mutagenesis is predicted (matching site/tissue or other). Brighter colors indicate the predictors where the epigenomic profile (CA or RT) matches the cancer type of the mutation profile. Asterisks indicate CA profiles of CD4-positive T-cells as predictors of SBS1 mutations. **B**. Prediction accuracy of megabase-scale burden of SBS signatures using CA and RT profiles. Signatures of carcinogens, unknown origin, and aging are more accurately predicted by CA and RT profiles than endogenous signatures. *P*-values are computed using F-tests with adjustment for genome-wide signature burden. Median accuracy values are printed. **C-F**. Top predictors of megabase-scale mutation burden in breast cancer quantified using SHAP scores. SHAP scores show the impact of a predictor (*i*.*e*., CA or RT profile) on the predictions (Y-axis) and corresponding predictor values (color gradient). **C**. Regional mutation burden in breast cancer genomes is predicted by CA profiles of primary breast cancers (BRCA) and uterine cancer (UCEC) as well as later replication timing (G2 phase) in a breast cancer cell line (MCF-7). CA profiles mostly negatively associate with mutagenesis while late RT profiles associate positively. **D**. Age-related mutations of SBS5 are predicted by BRCA CA profiles as well as RT profile of MCF-7, similarly to overall SNV burden. **E**. APOBEC-related mutations of SBS13 are also predicted by CA profiles (BRCA, as well as LUSC, TCGT); however, SHAP scores show that SBS13 mutations are positively correlated with CA. **F**. SBS1 mutations related to molecular clock activity are predicted by diverse CA profiles: two breast cancers as well as blood and immune cells (GM18520, CLL1_CD4, Peyer’s patch). Symbols indicate RT profiles (triangles) and CA profiles of normal tissues (circles).

SBS1 mutations showed the most variation in terms of CA profiles, compared to the profiles predictive of bulk mutations and other SBS signatures. Interestingly, the CA profile of CD4-positive T-cells from the peripheral blood of a CLL patient (CLL1_CD4) was consistently detected as a predictor of SBS1 mutation burden in six solid cancer types (liver, kidney, uterus, esophagus, stomach, head). This CA profile was only specific to SBS1 mutations and was not associated with bulk mutation burden or any other SBS signatures in the cancer types we studied. This T-cell CA profile may represent somatic mutations in non-cancerous cells of the immune system or the tumor microenvironment. As another example, SBS1 mutations in liver cancer associated with CA profiles of liver cancers, while overall regional mutation burden was predominantly associated with CA profiles of normal liver tissues. The clock-like SBS1 signature of 5-methylcytosine deamination is associated with cancer patient age and stem cell division rate, and this signature has been found in the somatic genomes of normal tissues and adult stem cells ^10,38,39^. Thus, SBS1 mutations may represent an earlier timepoint in tumor evolution or contribution from normal cells that remain convoluted in bulk tissue sequencing.

We asked if our random forest models were equally informative of various mutational signatures. Six classes of SBS signatures were compared in terms of prediction accuracy: APOBEC/AID, DNA-repair, carcinogens, two age-related classes (SBS1 and SBS5/40), and signatures of unknown cause, as predicted via all 869 CA and RT profiles in 17 cancer types. Three classes of signatures showed stronger associations with CA and RT profiles (**Figure 4B**): random forest predictions of carcinogenic signatures, signatures of unknown cause, and aging-associated signatures SBS5 and SBS40 were significantly more accurate than the DNA repair, APOBEC/AID, and SBS1 signatures combined, when accounting for number of mutations per signature as covariate of prediction accuracy (*P* ≤ 10^−4^; F-test) (**Supplementary Figure 4**). Thus, the mutational processes of carcinogen exposures, SBS5/40, and unknown signatures show stronger interactions with CA and RT in cancer genomes.

We studied the interactions of SBS signatures with CA and RT profiles in breast cancer (**Figure 4C-F**). SBS5 mutations, representing most mutations in the cohort, associated with four CA profiles of primary breast cancers (BRCA) and one late DNA-replicating profile of the breast cancer cell line MCF-7, reflecting tissue-specific associations of mutations and chromatin (**Figure 4D**). Regional mutation burden of all mutations was consistent with SBS5, and similarly associated with CA and RT profiles of breast cancer (**Figure 4C**). However, an additional predictive CA profile of uterine cancer was identified, perhaps due to common epigenetic features of female hormone-driven cancers. Megabase-scale burden of all mutations and the regional burden individual signatures negatively correlated with CA profiles in most cases, as expected. In contrast, SBS2 and SBS13 signatures of AID/APOBEC mutagenesis correlated positively with CA, such that higher SHAP values corresponded to increased chromatin accessibility (**Figure 4E**). This agrees with prior observations that AID targets epigenetically active elements and results in kataegis and clustered mutational signatures ^6,40,41^. Lastly, SBS1 mutations associated with three CA profiles representing peripheral blood, lymphoid follicles, and immune cells (**Figure 4F**), perhaps reflecting somatic mutagenesis in tumor-infiltrated immune cells and other cells in the tumor microenvironment. In summary, individual mutational signatures also predominantly associate with CA of primary cancers rather than normal tissues. The complex interactions of CA and RT with regional mutagenesis in certain mutational signatures may reflect inter-and intra-tumoral heterogeneity and help characterize the mechanisms of mutational processes.

### Excess mutations unexplained by epigenomes converge to cancer driver genes and developmental pathways

To quantify the regional mutagenesis unexplained by CA and RT, we investigated the genomic regions that were enriched in mutations relative to the mutation burden predicted by random forest regression. To enable a more detailed, gene-level functional analysis of enriched mutations, we repeated the predictions of regional mutation burden using a finer 100-kbps genomic resolution. This revealed 1570 unique genomic regions in 17 cancer types that were significantly enriched in mutations based on the CA-and RT-informed model residuals (*FDR* < 0.05) (**Figure 5A**). The mutation-enriched regions encoded 900 protein-coding genes including 67 known cancer genes ^42^, significantly more than expected by chance (33 expected, Fisher’s exact *P* = 3.1 × 10^−8^). Most driver genes were only found in single cancer types and represented key disease-specific drivers such as *EGFR* and *TERT* in glioma, *MYC* in BNHL and *APC* in colorectal cancer (**Supplementary Figure 5**). For example, in breast cancer, the region encoding *PIK3CA* was significantly enriched in mutations compared to the expected mutation burden based on the CA and RT landscape (92 SNVs observed *vs*. 44 expected; FDR = 9.2 × 10^−4^) (**Figure 5B**). PIK3CA is a major driver gene of breast cancer with hotspot mutations ^43^, thus showing that genome-wide statistical models of CA and RT can capture known driver genes.

**Figure 5.**
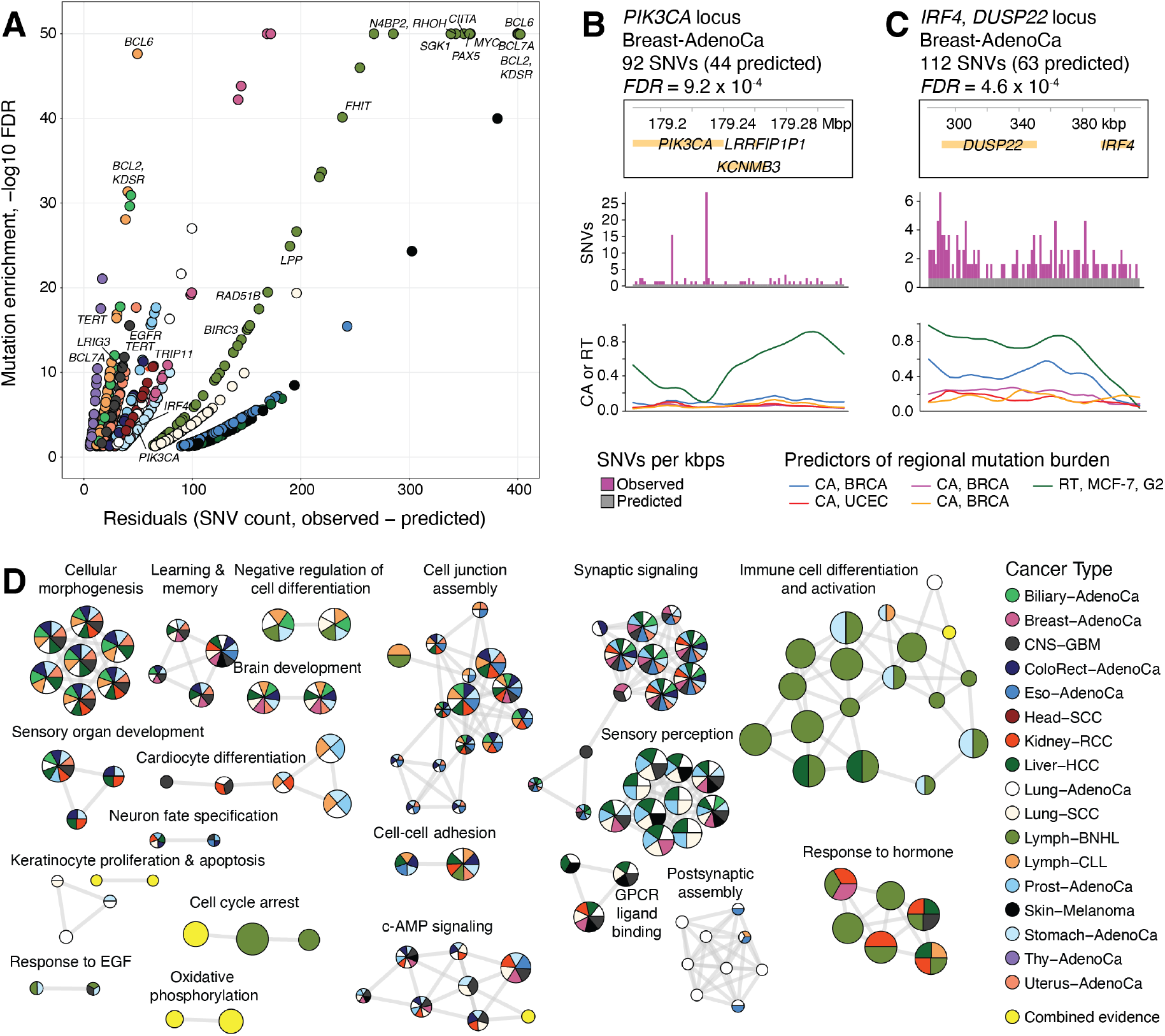
Excess mutation burden unexplained by CA and RT profiles converges to cancer genes and developmental pathways. **A**. Additional mutation burden exceeding predictions of CA and RT profiles, quantified via significantly higher model residuals of individual 100-kbps genomic windows. Scatter plot shows the genomic windows with elevated regional mutation burden (FDR < 0.05). Regions with known cancer genes are labeled if *FDR* < 10^−10^. Colors indicate cancer types (see panel D). X-axis is capped at 10^−50^ and Y-axis is capped at 400. **B-C**. Examples of genomic regions with excess mutation burden and known or putative cancer driver mutations. The plots show the genes in the region (top), mutation burden (SNVs per kbps; observed and expected) (middle), and the top five most significant predictors (*i*.*e*., CA and RT profiles; bottom). **B**. The genomic region encoding the driver gene *PIK3CA* in breast cancer. **C**. Super-enhancer region at the *IRF4*-*DUSP22* locus with elevated mutation burden in breast cancer. **D**. Pathway enrichment analysis of genomic regions with significantly higher regional mutation burden relative to CA and RT-informed predictions. The enrichment map shows significantly enriched biological processes and pathways (*FDR* < 0.05). The nodes represent enriched pathways and processes, the edges connect nodes sharing many genes, and the manually annotated subnetworks represent functionally related pathways and processes. Colors show the cancer types where the pathway enrichments were detected. Pathways and processes only detected in the joint analysis of multiple cancer types are shown in yellow.

The models also revealed regions with frequent non-coding mutations. The most prominent region was found in 12 of 17 cancer types due to unexpectedly frequent mutations, including breast cancer (112 SNVs observed *vs*. 63 expected; FDR = 4.6 × 10^−4^) (**Figure 5C; Supplementary Figure 6**). The region encodes the oncogenic transcriptional regulator *IRF4* (interferon regulatory factor 4) ^*44*^ and *DUSP22* encoding a signaling protein that was recently described as a network-implicated driver gene due to non-coding mutations ^45^. The region also includes super-enhancers of immune cells ^46^. The recurrence of mutations in this region in multiple cancer types highlights it as a potential pan-cancer region of interest.

We then asked whether the frequently-mutated regions were associated with common biological pathways and processes. An integrative pathway enrichment analysis that prioritized genomic regions detected in multiple cancer types revealed 177 significantly enriched pathways and processes (*FDR* < 0.05, ActivePathways ^47^), of which 142 (80%) were detected in more than one cancer type (**Figure 5D**). These findings converged into several functional themes of similar pathways and processes. First, developmental processes related to brain and the central nervous system, reproductive and sensory organs were associated with elevated mutation burden in multiple cancer types. Second, a group of processes related to synapse organization, olfactory and GPCR signaling were also identified in most cancer types. Third, cancer-related processes of cell cycle, hormone response, and signal transduction were also identified, often through pan-cancer data integration but not in any specific cancer type specifically. Lastly, a major group of processes related to immune system activation were predominantly detected in BNHL, potentially reflecting aberrant somatic hypermutation, as well as fewer associations with liver and stomach cancers.

This analysis shows that although individual frequently-mutated genomic regions are mostly characteristic of specific cancer types, enriched mutations converge to common pathways and processes in multiple cancer types. Convergence of these excess mutations to developmental and cancer-related processes is potentially explained by additional focal mutational processes targeting epigenetically active regions of the genome that are not captured by our models at the broader, sub-megabase resolution. Further, the enrichment of known cancer driver genes suggests that positive selection of functional mutations may also contribute to this additional mutation burden. This analysis exemplifies the complex interplay of cancer epigenomes, multi-scale mutational processes and positive selection of cancer genes.

## DISCUSSION

Our analysis highlights chromatin accessibility of primary human cancers as a major determinant of regional mutational processes in cancer genomes. Cancer epigenomes are predictive of regional mutation burden of matching cancer types, indicating tissue of origin associations in most cancer types we studied. While these associations are apparent for overall regional mutation burden in cancer genomes, they are also consistent with the regional variations in mutational signature burden. In contrast, the chromatin states of normal tissues and cell lines show only limited associations with regional mutagenesis of cancer genomes, extending the earlier studies that used the epigenetic profiles of cell lines and normal tissues to characterize mutational processes. The transformation of normal cells to cancer cells involves major changes in their epigenetic landscapes as gene-regulatory programs of cancer hallmark pathways are activated. Thus, one potential explanation to this stronger association of cancer epigenomes and regional mutagenesis is that mutational processes have a longer exposure on the somatic genomes shaped by the epigenomes of transformed cells, suggesting that many passenger mutations occur later in cancer evolution after the cells have acquired the epigenetic characteristics of cancer cells.

Replication timing information also associated with regional mutagenesis and confirmed strong effects with cell types related to cancer origin. However, CA profiles of primary human cancers evidently captured a larger fraction of variation of regional mutagenesis compared to RT profiles, apart from squamous cell and lymphoid cancers that strongly associated with relevant cell lines. Fewer RT profiles are used as predictors in our dataset and include mitotic cell lines that offer only limited representation of the diverse disease types in the pan-cancer cohort. As our models also include DNA replication timing profiles of several cell lines as reference, the stronger association with epigenomes of primary cancers shows that cancer epigenomes complement replication timing information with respect to regional mutagenesis. Interestingly, DNA replication has been shown to determine chromatin state ^48^. Thus, the informative CA profiles of human cancers may represent a proxy of cancer-specific replication dynamics.

Mutational signature analysis revealed interactions of mutational processes with CA and tissues of origin. Carcinogen signatures, as well as signatures of unknown etiology, were overall better predicted by CA and RT, in contrast to signatures of aging and DNA damage where the genome-wide predictions were less accurate. The stronger association of carcinogen signatures suggests that the chromatin environment interacts with DNA damage or repair processes of carcinogen exposure, for example through elevated mutational processes targeting active genes that are otherwise protected from mutations through error-free mismatch repair ^41^. Early replicating regions in cells exposed to tobacco mutagens show elevated mutagenesis in transcribed strands due to differential nucleotide excision repair activity ^49^. Based on their stronger interactions with RT and CA profiles, we speculate that some mutational signatures of currently unknown etiology may relate to carcinogens. SBS17 mutations show some of the strongest interactions with CA and RT in stomach and esophageal cancers in our analysis. This signature is currently of unknown cause, however it has been linked to gastric acid reflux and reactive oxygen species ^24^. Further integrative analysis of clinical and lifestyle information with patterns of regional mutagenesis may shed light to these mutational processes.

We observed a functional convergence to developmental processes and cancer-related pathways in the genomic regions where mutations were enriched beyond the predictions of our epigenomic models. On the one hand, these data suggest that additional mutational processes affect distinct regions with developmental genes and open-chromatin regions in individual cancer types, however these regions converge to the same molecular pathways across cancer types. Such local mutational processes are consistent with previous studies. For example, transcription start sites of highly expressed genes and constitutively-bound binding sites of CTCF are subject to elevated local mutagenesis in multiple cancer types ^16^. Lineage-specific genes are enriched in indel mutations in solid cancers ^50^. Such local mutational processes are complementary to megabase-scale processes where open chromatin is generally associated with a lower mutation frequency. On the other hand, the enrichment of cancer genes and pathways in our data suggests that some mutations unexplained by CA and RT are functional in cancer and their frequent occurrence at specific genes, non-coding elements and molecular pathways is explained by positive selection ^3-5,45^. We can use this computational framework to find genomic regions with known and putative driver mutations in coding and non-coding sites. Further study of these regions may deepen our understanding of mutational processes and refine the catalogues of driver mutations.

This approach enables future studies to decipher the mechanisms and phenotypic associations of mutational processes. Clinical, genetic, and epigenetic profiles of cancer patients can be integrated to understand how regional mutational processes and the chromatin landscape are modulated by clinical variables such as stage, grade or the therapies applied, genetic features such as somatic driver mutations or inherited cancer risk variants, or lifestyle choices such as tobacco or alcohol consumption. Complementary insights from sub-clonal reconstruction analysis of cancer genomes ^2,51^, as well as single-cell sequencing of genomes and epigenomes will allow mapping of regional mutagenesis at the level of distinct cell populations contributing to temporal and spatial variation in mutational processes. As such multimodal datasets grow, we can learn about early cancer evolution by comparing regional mutagenesis in the genomes of cancers and normal cells. Understanding the molecular and genetic determinants of regional mutagenesis and signatures in cancer genomes may help characterize carcinogen exposures and genetic predisposition, ultimately enhancing early cancer detection and prevention in the future.

## Methods

### Somatic mutations in whole cancer genomes

We analyzed somatic single nucleotide variants (SNVs; n = 43,778,859) derived from whole-genome sequencing (WGS) of 2,583 primary cancer samples that were uniformly mapped to GRCh37/hg19 as part of the ICGC/TCGA Pan-cancer Analysis of Whole Genomes (PCAWG) project ^1^. Indels and variants in sex chromosomes were excluded. To integrate mutations with epigenetic information, we mapped the SNVs to the human genome version GRCh38 using the LiftOver function of the rtracklayer package in R (v 1.48) ^52^. We removed 66 hypermutated tumors with more than 90,000 mutations (∼30 mutations / Mbps), resulting in a dataset of 23,215,600 SNVs in 2,517 whole cancer genomes. We analyzed the genomes of 17 cancer types with at least 25 samples in PCAWG as well as related epigenetic profiles of normal and cancer tissues, as well as the pan-cancer dataset of all 37 cancer types.

### Chromatin accessibility (CA) and replication timing (RT) profiles

Chromatin accessibility data was derived from several ATAC-seq datasets, including the ENCODE3 project and six additional studies to maximize the coverage of cancer types included in the PCAWG dataset, as described below. CA profiles of 196 human cell and tissue types and 9 cancer cell lines at a single basepair (bp) resolution in GRCh38 were derived from the ENCODE3 project ^27^. CA profiles of 115 normal human brain samples at a single bp resolution in GRCh37 were retrieved from the study by Fullard *et al*. ^29^. CA profiles of 21 normal immune cell types (B-cells, T-cells) and 34 primary cancers (CLL) at a single base pair resolution in GRCh37 were retrieved from the study by Rendeiro *et al*. ^30^. Four CA profiles of HEK293 embryonic kidney cells at a 10-bp resolution in GRCh37 were retrieved from the study by Karabacak Calviello *et al*. ^31^. CA profiles for two lymphoma cell lines at a single-bp resolution in GRCh37 were retrieved from the study by Scharer *et al*. ^32^. One CA profile of the normal human melanocytes (NHM1) cell line at a 10-bp resolution in GRCh37 were retrieved from the study by Fontanals-Cirera *et al*. ^33^. CA profiles of four normal prostate tissues and six primary prostate cancers at a single-bp resolution in GRCh37 were retrieved from the study by Pomerantz *et al*. ^34^. CA profiles of several cancer types were retrieved from the TCGA ATAC-seq dataset ^28^ of 410 primary cancer samples, representing cancers of 404 unique patient donors and 796 genome-wide profiles in total. We used 381 CA profiles of the TCGA dataset such that technical and biological replicates of distinct cancer samples were pooled by per-region averaging CA signal. Prior to this averaging, 22 CA profiles for which only one replicate was available were removed, and one CA profile of a low-grade glioma (LGG) that was an outlier in our initial analyses as also removed. In total, 773 CA profiles were included in the analysis, including 421 cancer profiles, 341 profiles of normal tissues and cell lines, and 11 profiles of cancer cell lines. Besides these CA profiles, 96 replication timing profiles of 16 cell lines, each with six cell cycle phases, were derived from the RepliSeq study by Hansen *et al*. ^35^. CA and RT profiles were constructed from the BigWig files of the original studies using mean values of signal intensity per each genomic window. Genomic coordinates of the GRCh37 reference genome were mapped to the GRCh38 reference genome using LiftOver. In total, the set of 869 (773 + 96) CA and RT profiles was used.

### Integrating regional mutagenesis with CA and RT profiles

We evaluated chromatin accessibility, replication timing and mutation burden in non-overlapping genomic regions of one megabase (Mbp; one million base pairs). We excluded a subset of genomic regions with low mappability (≤80% in the UMAP software ^53^) as well as sex chromosomes, resulting in 2,465 regions included in the study. For megabase-scale CA and RT profiles, each genomic region was assigned the mean value of its epigenetic signal. For megabase-scale somatic mutation burden, each region was assigned the total mutation count separately for the pan-cancer dataset and each of the 17 cancer types. In two cohorts (chronic lymphocytic leukemia; B-cell non-Hodgkin lymphoma), we removed two regions encoding immunoglobulin genes (chr2:89Mbps, chr22:23Mbps) with known high somatic variation in immune cells, as observed in our initial analyses.

### Random forest regression

Megabase-scale profiles of mutation burden and CA and RT profiles were analyzed with random forest regression ^54^ with CA and RT profiles as the predictors (*i*.*e*., features) and mutation burden as the target (*i*.*e*., response). Number of trees (1000) and fraction of predictors at each split (1/3) were used as hyperparameters. Monte-Carlo cross-validation over 1000 data splits considered subsets of genomic regions for model training (80%) and validation (20%). We used the adjusted R^2^ (adj.R^2^) metric to evaluate model performance that measures the variance explained by the model adjusted for model complexity (*i*.*e*., the number of CA and RT profiles used for predictions).

### Comparing CA profiles of primary cancers and normal tissues as predictors of regional mutagenesis

First, we compared the overall accuracy of predicting megabase-scale mutation burden using CA profiles of cancers *vs*. normal tissues. Two sets of random forest regression models were run in a joint Monte-Carlo cross-validation procedure that used all CA profiles of normal tissues (M_n_) and cancers (M_c_) as predictors, respectively. Both models also included the same set of RT profiles as predictors as reference. At each iteration, models were trained on matching subsets of genomic regions (80%) and tested on the remaining genomic regions (20%), and model accuracy (adj.R^2^) values as well as the relative change values (Δadj.R^2^ = adj.R^2^(M_c_)-adj.R^2^(M_n_)) were derived in the corresponding test sets, allowing us to directly compare the two models. For each cancer type, median Δadj.R^2^ values and 95% confidence intervals were reported. Empirical P-values were computed as the fraction of cross-validation iterations where the Δadj.R^2^ crossed on the opposite side of zero relative to the median value. We also trained the models M_n_ and M_c_ on the full set of genomic regions and compared the accuracy of the two sets of models. Observed and model-predicted mutation burden values per region were visualized as scatter plots with local regression (loess) trendlines (span=0.9). Spearman correlation tests were used to evaluate the associations of model accuracy, WGS cohort size and per-megabase mutation burden in different cancer types.

### Evaluating CA and RT profiles as predictors of regional mutation burden

We used the incMSE (increase in model mean-squared-error) metric to evaluate the most important features (*i*.*e*., CA, RT profiles) in random forest models. incMSE measures the relative change in model prediction accuracy upon permutations of the values of a given feature. We derived incMSE values of CA and RT profiles for the 17 cancer types for which matching CA and/or RT profiles were available. Two additional statistical methods were used to evaluate the significance of incMSE of CA and RT profiles. First, permutation tests were used to detect CA and RT profiles where incMSE values significantly exceeded those of permuted data. We fitted random forest regression models for every cancer type 1,000 times using randomly reassigned megabase-scale mutation burden estimates as null distributions for incMSE values for CA and RT profiles. Specific profiles were considered statistically significant if their observed incMSE values exceeded all 1000 incMSE values from permuted datasets (*i*.*e*., empirical *P* < 0.001). Second, we used bootstrapping of random forest regression where the genomic regions with predictor and response values were sampled randomly with replacement. We repeated this resampling process 1000 times and recorded the incMSE values for all CA and RT profiles to evaluate the confidence intervals of the derived incMSEs.

### Feature importance of CA and RT profiles in predictions of regional mutagenesis

We used the SHapley Additive exPlanation (SHAP) method ^36,55^ to interpret the interactions of regional mutation burden and CA and RT profiles. Here, SHAP scores reflect the importance of each feature in the random forest model (*i*.*e*., CA or RT profile) in predicting a specific observation (*i*.*e*., mutation burden of a certain genomic region), and represent its relative contribution to the prediction (*i*.*e*., effect size) as well as the direction of the prediction (*i*.*e*., positive or negative). SHAP values were computed using models trained all genomic regions and separately for cancer types, using the python packages shap (0.35.0) ^36,55^ and scikit-learn (0.23.1) ^56^ via the R package reticulate (1.16) ^57^.

### Associating mutational signatures with CA and RT

For mutational signature analysis, we used single base substitution (SBS) annotations of SNVs derived in the PCAWG project ^6^. For each genomic region, we computed the mutational signature burden probabilistically by adding the SBS-specific probabilities of all individual SNVs in the region, thus accounting for all signature exposures rather than top-ranking signatures for each SNV. We filtered lower-frequency SBS signatures in each cancer type (*i*.*e*., <20,000 or <5% of all SNVs). To evaluate CA and RT profiles as predictors of megabase-scale mutational signature burden, we trained random forest models where a probabilistic SBS profile was used as model response. We evaluated model performance, selected top features, and computed SHAP scores similarly to bulk mutation analysis described above. We grouped the mutational signatures based on their etiology according to the COSMIC database (version 3.2, downloaded in March 2021): AID/APOBEC, deficient DNA repair, exogeneous/carcinogen, unknown/other, SBS5/40 and SBS1. We compared model accuracy values for predicting regional mutational signature burden of the six classes of signatures using ANOVA analysis and F-tests. We used the covariate of the average megabase-scale SBS burden to account for a potential of improved predictions in cancer types with higher overall mutation burden.

### Prioritizing highly-mutated genomic regions exceeding CA and RT predictions

To study regional mutation burden unexplained by CA and RT profiles, we prioritized the genomic regions where the random forest predictions significantly underestimated the observed mutation burden. Random forest regression was repeated on 100-kbps regions to improve gene-level interpretation. To score genomic regions, we subtracted the predicted mutation counts from the observed counts to derive residual values. Residuals were Z-transformed and the resulting one-tailed *P*-values were adjusted for multiple testing using Benjamini-Hochberg FDR.

### Pathway enrichment analysis of regional mutation variation

To understand the functional importance of excess mutations unexplained by CA and RT profiles, we performed an integrative pathway enrichment analysis across the relevant cancer types using the ActivePathways method ^47^ (*FDR* < 0.05). Gene sets of biological processes of Gene Ontology and molecular pathways of Reactome were collected from the GMT files provided in the g:Profiler web server ^58^ (downloaded Feb 23^rd^, 2021) and were filtered using default settings of ActivePathways. In each cancer type, all protein-coding genes were assigned the P-values reflecting excess mutation burden unexplained by CA and RT in respective regions. The data fusion in ActivePathways prioritized the genes that were frequently mutated in multiple cancer types. Enriched pathways were visualized as an enrichment map ^59^ and themes were curated manually. We also visualized the genomic regions with excess mutations as a scatter plot of residual values and -log10-transformed FDR values that were capped at 400, and 10^−50^, respectively. We highlighted known cancer driver genes of the Cancer Gene Census database ^42^ (downloaded Mar 26^th^ 2021) and computed their enrichment in the list of pathway-associated genes using a Fisher’s exact test.

## Code availability

Source code for this study is available at https://github.com/reimandlab/CA2M_v2.

## Acknowledgments

We thank Christian A. Lee, Kevin Cheng, Phedias Diamandis, and Anne Martel for constructive comments on this study. This work was supported by the Canadian Institutes of Health Research (CIHR) Project Grant to J.R., A New Investigator Award of the Terry Fox Research Institute (TFRI) to J.R., and the Investigator Award to J.R. from the Ontario Institute for Cancer Research (OICR). Funding to OICR is provided by the Government of Ontario. The results shown here are in whole or part based upon data generated by the TCGA Research Network: https://www.cancer.gov/tcga. We acknowledge the contributions of the many clinical networks of ICGC and TCGA who provided samples and data to PCAWG. We thank the patients and their families for their participation in ICGC and TCGA projects.

## Author contributions

O.O. analyzed the data and prepared the figures. J.R. and O.O. interpreted the data and wrote the manuscript. J.R. conceived and supervised the project. The authors reviewed and edited the manuscript and approved the final version.

## References

1 ICGC-TCGA Pan-Cancer Analysis of Whole Genomes Consortium. Pan-cancer analysis of whole genomes. Nature 578, 82–93, doi:10.1038/s41586-020-1969-6 (2020).

2 Gerstung, M. et al. The evolutionary history of 2,658 cancers. Nature 578, 122–128, doi:10.1038/s41586-019-1907-7 (2020).

3 Martincorena, I. et al. Universal Patterns of Selection in Cancer and Somatic Tissues. Cell 171, 1029–1041 e1021, doi:10.1016/j.cell.2017.09.042 (2017).

4 Rheinbay, E. et al. Analyses of non-coding somatic drivers in 2,693 cancer whole genomes. Nature 578, 102–111 (2020).

5 Zhu, H. et al. Candidate Cancer Driver Mutations in Distal Regulatory Elements and Long-Range Chromatin Interaction Networks. Mol Cell, doi:10.1016/j.molcel.2019.12.027 (2020).

6 Alexandrov, L. B. et al. The repertoire of mutational signatures in human cancer. Nature 578, 94–101, doi:10.1038/s41586-020-1943-3 (2020).

7 Li, Y. et al. Patterns of somatic structural variation in human cancer genomes. Nature 578, 112–121, doi:10.1038/s41586-019-1913-9 (2020).

8 Kumar, S. et al. Passenger Mutations in More Than 2,500 Cancer Genomes: Overall Molecular Functional Impact and Consequences. Cell 180, 915–927 e916, doi:10.1016/j.cell.2020.01.032 (2020).

9 Martincorena, I. et al. Tumor evolution. High burden and pervasive positive selection of somatic mutations in normal human skin. Science 348, 880–886, doi:10.1126/science.aaa6806 (2015).

10 Blokzijl, F. et al. Tissue-specific mutation accumulation in human adult stem cells during life. Nature 538, 260–264, doi:10.1038/nature19768 (2016).

11 Supek, F. & Lehner, B. Scales and mechanisms of somatic mutation rate variation across the human genome. DNA Repair (Amst), 102647, doi:10.1016/j.dnarep.2019.102647 (2019).

12 Gonzalez-Perez, A., Sabarinathan, R. & Lopez-Bigas, N. Local Determinants of the Mutational Landscape of the Human Genome. Cell 177, 101–114, doi:10.1016/j.cell.2019.02.051 (2019).

13 Pich, O. et al. The mutational footprints of cancer therapies. Nature genetics 51, 1732–1740, doi:10.1038/s41588-019-0525-5 (2019).

14 Katainen, R. et al. CTCF/cohesin-binding sites are frequently mutated in cancer. Nature genetics 47, 818–821, doi:10.1038/ng.3335 (2015).

15 Sabarinathan, R., Mularoni, L., Deu-Pons, J., Gonzalez-Perez, A. & Lopez-Bigas, N. Nucleotide excision repair is impaired by binding of transcription factors to DNA. Nature 532, 264–267, doi:10.1038/nature17661 (2016).

16 Lee, C. A., Abd-Rabbo, D. & Reimand, J. Functional and genetic determinants of mutation rate variability in regulatory elements of cancer genomes. Genome Biol 22, 133, doi:10.1186/s13059-021-02318-x (2021).

17 Lawrence, M. S. et al. Mutational heterogeneity in cancer and the search for new cancer-associated genes. Nature 499, 214–218, doi:10.1038/nature12213 (2013).

18 Schuster-Bockler, B. & Lehner, B. Chromatin organization is a major influence on regional mutation rates in human cancer cells. Nature 488, 504–507, doi:10.1038/nature11273 (2012).

19 Stamatoyannopoulos, J. A. et al. Human mutation rate associated with DNA replication timing. Nature genetics 41, 393–395, doi:10.1038/ng.363 (2009).

20 Supek, F. & Lehner, B. Differential DNA mismatch repair underlies mutation rate variation across the human genome. Nature 521, 81–84, doi:10.1038/nature14173 (2015).

21 Zheng, C. L. et al. Transcription restores DNA repair to heterochromatin, determining regional mutation rates in cancer genomes. Cell Rep 9, 1228–1234, doi:10.1016/j.celrep.2014.10.031 (2014).

22 Woo, Y. H. & Li, W. H. DNA replication timing and selection shape the landscape of nucleotide variation in cancer genomes. Nat Commun 3, 1004, doi:10.1038/ncomms1982 (2012).

23 Liu, L., De, S. & Michor, F. DNA replication timing and higher-order nuclear organization determine single-nucleotide substitution patterns in cancer genomes. Nat Commun 4, 1502, doi:10.1038/ncomms2502 (2013).

24 Tomkova, M., Tomek, J., Kriaucionis, S. & Schuster-Bockler, B. Mutational signature distribution varies with DNA replication timing and strand asymmetry. Genome Biol 19, 129, doi:10.1186/s13059-018-1509-y (2018).

25 Jiao, W. et al. A deep learning system accurately classifies primary and metastatic cancers using passenger mutation patterns. Nat Commun 11, 728, doi:10.1038/s41467-019-13825-8 (2020).

26 Polak, P. et al. Cell-of-origin chromatin organization shapes the mutational landscape of cancer. Nature 518, 360–364, doi:10.1038/nature14221 (2015).

27 Consortium, E. P. et al. Expanded encyclopaedias of DNA elements in the human and mouse genomes. Nature 583, 699–710, doi:10.1038/s41586-020-2493-4 (2020).

28 Corces, M. R. et al. The chromatin accessibility landscape of primary human cancers. Science 362, doi:10.1126/science.aav1898 (2018).

29 Fullard, J. F. et al. An atlas of chromatin accessibility in the adult human brain. Genome research 28, 1243–1252, doi:10.1101/gr.232488.117 (2018).

30 Rendeiro, A. F. et al. Chromatin mapping and single-cell immune profiling define the temporal dynamics of ibrutinib response in CLL. Nat Commun 11, 577, doi:10.1038/s41467-019-14081-6 (2020).

31 Karabacak Calviello, A., Hirsekorn, A., Wurmus, R., Yusuf, D. & Ohler, U. Reproducible inference of transcription factor footprints in ATAC-seq and DNase-seq datasets using protocol-specific bias modeling. Genome Biol 20, 42, doi:10.1186/s13059-019-1654-y (2019).

32 Scharer, C. D. et al. Genome-wide CIITA-binding profile identifies sequence preferences that dictate function versus recruitment. Nucleic Acids Res 43, 3128–3142, doi:10.1093/nar/gkv182 (2015).

33 Fontanals-Cirera, B. et al. Harnessing BET Inhibitor Sensitivity Reveals AMIGO2 as a Melanoma Survival Gene. Mol Cell 68, 731–744 e739, doi:10.1016/j.molcel.2017.11.004 (2017).

34 Pomerantz, M. M. et al. Prostate cancer reactivates developmental epigenomic programs during metastatic progression. Nature genetics 52, 790–799, doi:10.1038/s41588-020-0664-8 (2020).

35 Hansen, R. S. et al. Sequencing newly replicated DNA reveals widespread plasticity in human replication timing. Proc Natl Acad Sci U S A 107, 139–144, doi:10.1073/pnas.0912402107 (2010).

36 Lundberg, S. M. et al. From Local Explanations to Global Understanding with Explainable AI for Trees. Nat Mach Intell 2, 56–67, doi:10.1038/s42256-019-0138-9 (2020).

37 Odegard, V. H. & Schatz, D. G. Targeting of somatic hypermutation. Nat Rev Immunol 6, 573–583, doi:10.1038/nri1896 (2006).

38 Alexandrov, L. B. et al. Clock-like mutational processes in human somatic cells. Nature genetics 47, 1402–1407, doi:10.1038/ng.3441 (2015).

39 Abascal, F. et al. Somatic mutation landscapes at single-molecule resolution. Nature, doi:10.1038/s41586-021-03477-4 (2021).

40 Wang, Q. et al. Epigenetic targeting of activation-induced cytidine deaminase. Proc Natl Acad Sci U S A 111, 18667–18672, doi:10.1073/pnas.1420575111 (2014).

41 Supek, F. & Lehner, B. Clustered Mutation Signatures Reveal that Error-Prone DNA Repair Targets Mutations to Active Genes. Cell 170, 534–547 e523, doi:10.1016/j.cell.2017.07.003 (2017).

42 Futreal, P. A. et al. A census of human cancer genes. Nat Rev Cancer 4, 177–183, doi:10.1038/nrc1299 (2004).

43 Nik-Zainal, S. et al. Landscape of somatic mutations in 560 breast cancer whole-genome sequences. Nature 534, 47–54, doi:10.1038/nature17676 (2016).

44 Iida, S. et al. Deregulation of MUM1/IRF4 by chromosomal translocation in multiple myeloma. Nature genetics 17, 226–230, doi:10.1038/ng1097-226 (1997).

45 Reyna, M. A. et al. Pathway and network analysis of more than 2,500 whole cancer genomes. Nature Communications 11, 729 (2020).

46 Hnisz, D. et al. Super-enhancers in the control of cell identity and disease. Cell 155, 934–947, doi:10.1016/j.cell.2013.09.053 (2013).

47 Paczkowska, M. et al. Integrative pathway enrichment analysis of multivariate omics data. Nature Communications 11, 735 (2020).

48 Klein, K. N. et al. Replication timing maintains the global epigenetic state in human cells. Science 372, 371–378, doi:10.1126/science.aba5545 (2021).

49 Kucab, J. E. et al. A Compendium of Mutational Signatures of Environmental Agents. Cell 177, 821–836 e816, doi:10.1016/j.cell.2019.03.001 (2019).

50 Imielinski, M., Guo, G. & Meyerson, M. Insertions and Deletions Target Lineage-Defining Genes in Human Cancers. Cell 168, 460–472 e414, doi:10.1016/j.cell.2016.12.025 (2017).

51 Dentro, S. C. et al. Characterizing genetic intra-tumor heterogeneity across 2,658 human cancer genomes. Cell, doi:10.1016/j.cell.2021.03.009 (2021).

52 Lawrence, M., Gentleman, R. & Carey, V. rtracklayer: an R package for interfacing with genome browsers. Bioinformatics 25, 1841–1842, doi:10.1093/bioinformatics/btp328 (2009).

53 Karimzadeh, M., Ernst, C., Kundaje, A. & Hoffman, M. M. Umap and Bismap: quantifying genome and methylome mappability. Nucleic Acids Res 46, e120, doi:10.1093/nar/gky677 (2018).

54 Ho, T. K. Random decision forests. IEEE Proceedings of 3rd international conference on document analysis and recognition 1, 278–282, doi:doi:10.1109/ICDAR.1995.598994 (1995).

55 Lundberg, S. M. & Lee, S. A Unified Approach to Interpreting Model Predictions. Advances in Neural Information Processing Systems (NIPS) 30 (2017).

56 Pedregosa, F. et al. Scikit-learn: Machine Learning in Python. Journal of Machine Learning Research 12, 2825–2830 (2011).

57 Ushey, K., Allaire, J. J. & Tang, Y. Reticulate R Package. GitHub https://rstudio.github.io/reticulate/ doi:https://rstudio.github.io/reticulate/ (2021).

58 Reimand, J., Kull, M., Peterson, H., Hansen, J. & Vilo, J. g:Profiler--a web-based toolset for functional profiling of gene lists from large-scale experiments. Nucleic acids research 35, W193–200, doi:10.1093/nar/gkm226 (2007).

59 Reimand, J. et al. Pathway enrichment analysis and visualization of omics data using g:Profiler, GSEA, Cytoscape and EnrichmentMap. Nat Protoc 14, 482–517, doi:10.1038/s41596-018-0103-9 (2019).

