## Supplementary Figures for "Chromatin accessibility of primary human cancers ties regional mutational processes with tissues of origin"

Supplementary information
Ocsenas and Reimand, 2021

| Study Name | Project Code | Number of samples | Number of mutations | Included separately |
|---|---|---|---|---|
| Liver Hepatocellular carcinoma | Liver–HCC | 314 | 3764099 | Included |
| Pancreas Adenocarcinoma | Panc–AdenoCa | 232 | 1492798 | Excluded |
| Prostate adenocarcinoma | Prost–AdenoCa | 199 | 637414 | Included |
| Breast Adenocarcinoma | Breast–AdenoCa | 195 | 1396733 | Included |
| Kidney Renal cell carcinoma | Kidney–RCC | 143 | 897708 | Included |
| Medulloblastoma | CNS–Medullo | 141 | 199596 | Excluded |
| Ovary Adenocarcinoma | Ovary–AdenoCa | 110 | 965271 | Excluded |
| Diffuse large B–cell lymphoma | Lymph–BNHL | 104 | 1130214 | Included |
| Esophageal Adenocarcinoma | Eso–AdenoCa | 95 | 2465375 | Included |
| Chronic lymphocytic leukemia | Lymph–CLL | 90 | 217704 | Included |
| Pilocytic Astrocytoma | CNS–PiloAstro | 89 | 22005 | Excluded |
| Pancreatic Neuroendocrine tumour | Panc–Endocrine | 81 | 252855 | Excluded |
| Stomach Adenocarcinoma | Stomach–AdenoCa | 66 | 1048100 | Included |
| Melanoma | Skin–Melanoma | 65 | 2274735 | Included |
| Head/Neck Squamous cell carcinoma | Head–SCC | 56 | 876369 | Included |
| Thyroid Adenocarcinoma | Thy–AdenoCa | 48 | 64943 | Included |
| Lung Squamous cell carinoma | Lung–SCC | 45 | 1824949 | Included |
| Kidney Renal cell carcinoma, chromophobe type | Kidney–ChRCC | 43 | 78318 | Excluded |
| Colorectal Adenocarcinoma | ColoRect–AdenoCa | 43 | 664599 | Included |
| Uterus Adenocarcinoma | Uterus–AdenoCa | 42 | 487894 | Included |
| Bone Osteosarcoma | Bone–Osteosarc | 41 | 157718 | Excluded |
| Glioblastoma multiforme | CNS–GBM | 38 | 263689 | Included |
| Bone Leiomyosarcoma | Bone–Leiomyo | 34 | 183141 | Excluded |
| Biliary Adenocarcinoma | Biliary–AdenoCa | 33 | 243049 | Included |
| Lung Adenocarcinoma | Lung–AdenoCa | 33 | 725328 | Included |
| Bladder Transitional cell carcinoma | Bladder–TCC | 23 | 504886 | Excluded |
| Myeloid Myeloproliferative neoplasm | Myeloid–MPN | 23 | 24171 | Excluded |
| Oligodendroglioma | CNS–Oligo | 18 | 48178 | Excluded |
| Cervix Squamous cell carcinoma | Cervix–SCC | 18 | 114186 | Excluded |
| Myeloid Acute myeloid leukemia | Myeloid–AML | 13 | 19265 | Excluded |
| Breast Lobular carcinoma | Breast–LobularCa | 13 | 96983 | Excluded |
| Bone neoplasm, epithelioid | Bone–Epith | 11 | 23424 | Excluded |
| Chondroblastoma | Bone–Cart | 9 | 7846 | Excluded |
| Breast In situ adenocarcinoma | Breast–DCIS | 3 | 6012 | Excluded |
| Lymphoid (Not otherwise specified) | Lymph–NOS | 2 | 26272 | Excluded |
| Myelodysplastic syndrome | Myeloid–MDS | 2 | 1624 | Excluded |
| Cervix Adenocarcinoma | Cervix–AdenoCa | 2 | 8149 | Excluded |
| TOTAL | PANCAN | 2517 | 23215600 | |

**Supplementary Figure 1.** Summary of whole cancer genome datasets of the PCAWG project included in the analysis. The rightmost column indicates whether the cancer type was included separately for regional mutation burden analysis. All cancer were included in the pan-cancer analysis.

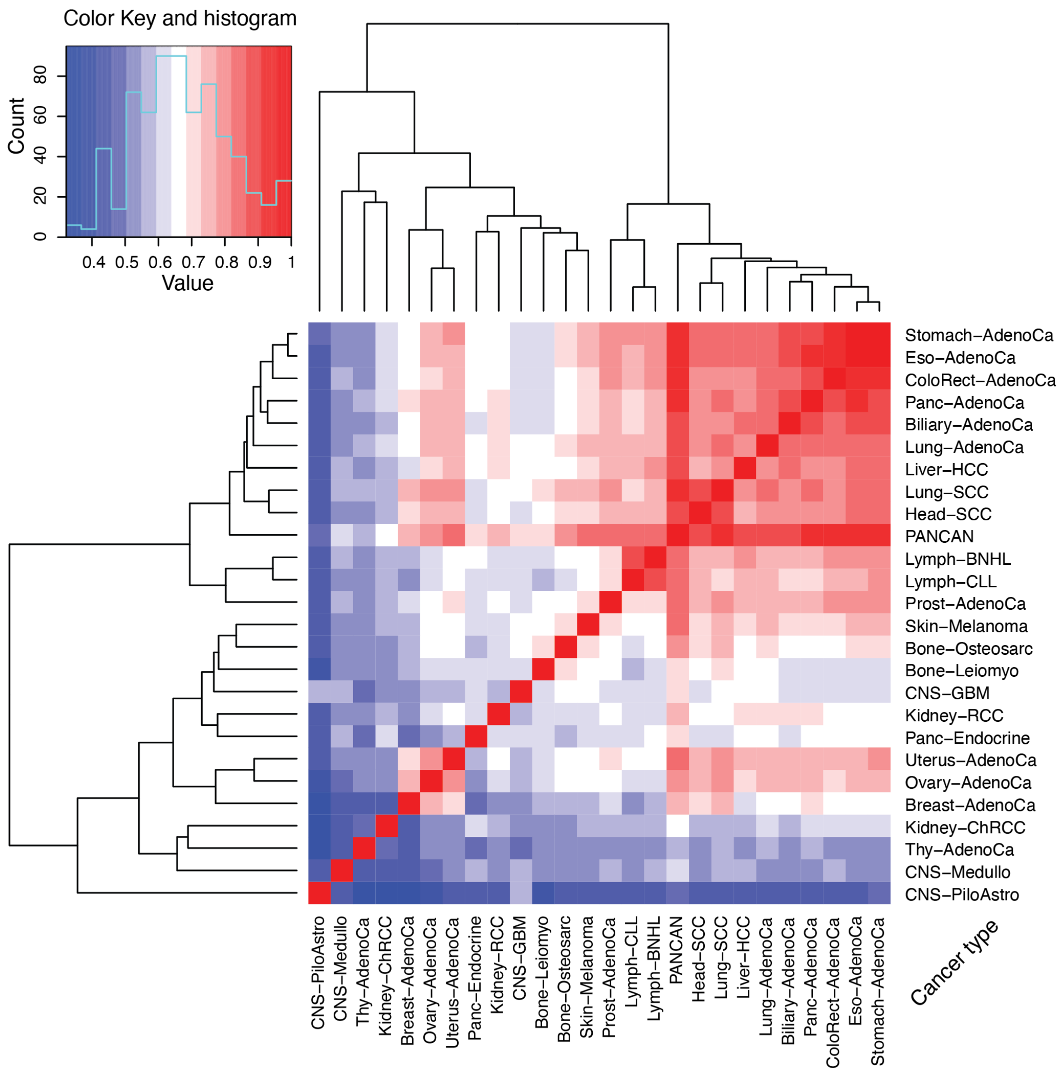

**Supplementary Figure 2.** Correlation heatmap of megabase-scale mutation burden in cancer types in the PCAWG datasets. Clustering suggests similarity of cancer types in similar anatomical sites and cells of origin. For example, digestive tract cancers (stomach, esophageal, colorectal), as well as squamous cell cancers (lung, head & neck) are clustered.

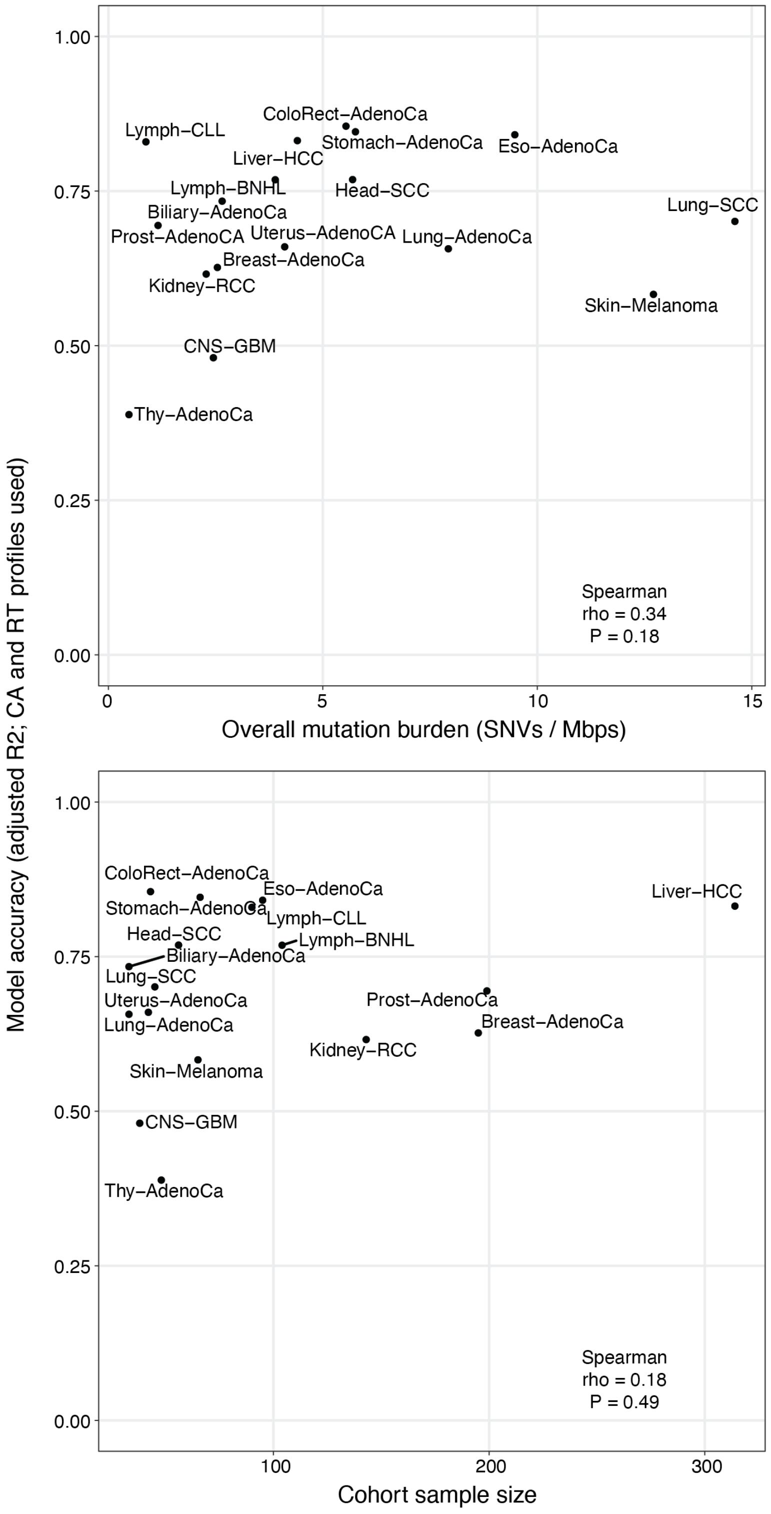

**Supplementary Figure 3.** Model accuracy of megabase-scale mutation burden prediction shows limited correlations with overall mutation burden (top) and cohort size (bottom).

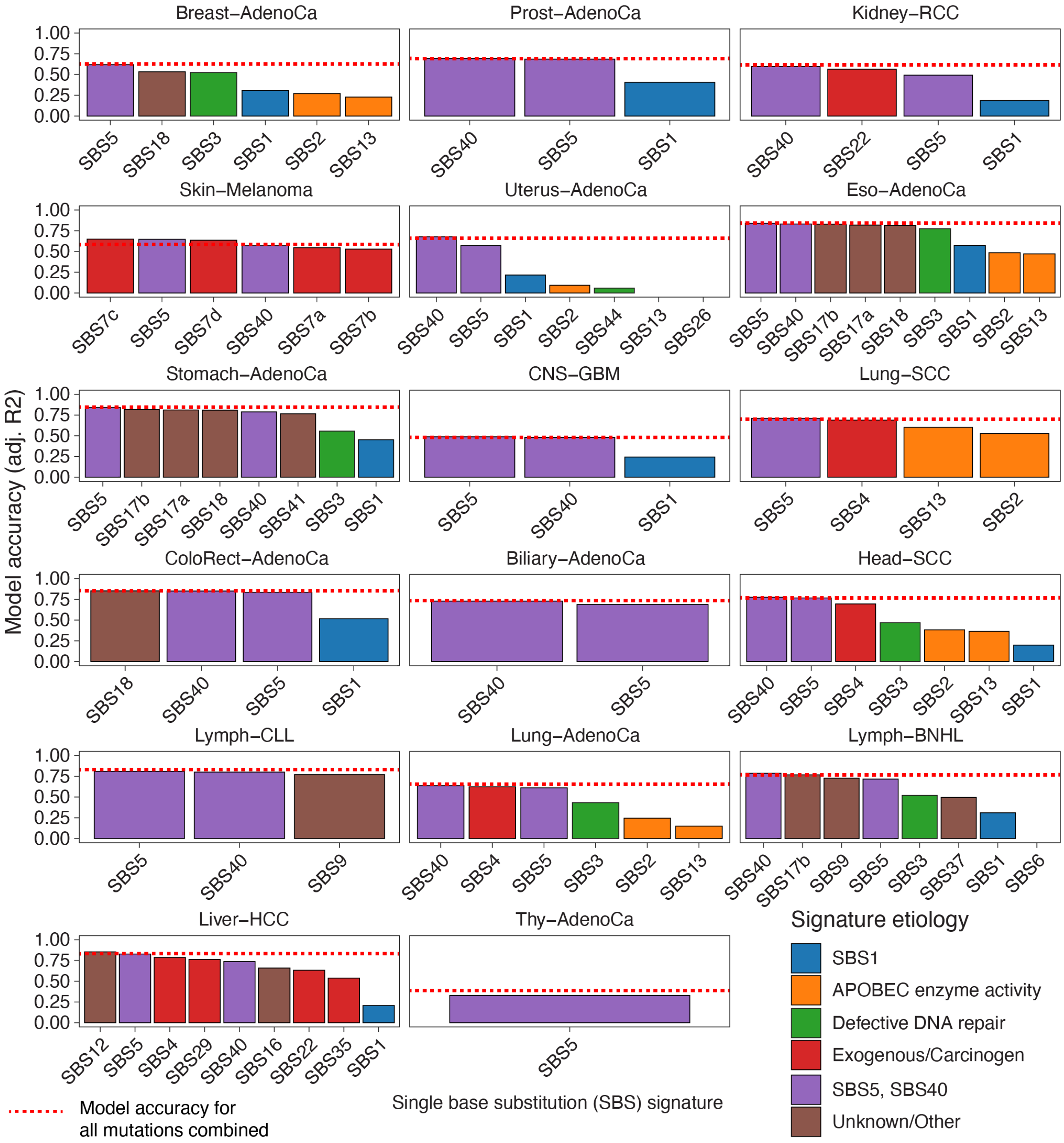

**Supplementary Figure 4.** Prediction accuracy of regional mutation burden of SNVs associated with specific single base substitution (SBS) signatures.

| Gene | Cancer types with mutations | CGC cancer types | CGC role |
|---|---|---|---|
| AFF3 | Breast–AdenoCa | ALL, T–ALL | oncogene, fusion |
| APC | ColoRect–AdenoCa | colorectal, pancreatic, desmoid, hepatoblastoma, glioma, other CNS | TSG |
| BCL2 | Lymph–BNHL | NHL, CLL | oncogene, fusion |
| BCL6 | Lymph–CLL, Lymph–BNHL | NHL, CLL | oncogene, fusion |
| BCL7A | Lymph–CLL, Lymph–BNHL | BNHL | fusion |
| BIRC3 | Lymph–BNHL | CLL, MALT, MCL, MM | oncogene, TSG, fusion |
| CASP8 | Head–SCC | hepatocellular, oral squamous cell, breast | TSG |
| CCND1 | Head–SCC | CLL, B–ALL, breast | oncogene, fusion |
| CD79A | Thy–AdenoCa | DLBCL, WM | oncogene |
| CD79B | Breast–AdenoCa | DLBCL, WM | oncogene |
| CDH10 | Lung–SCC | melanoma | TSG |
| CDH11 | ColoRect–AdenoCa | aneurysmal bone cyst | TSG, fusion |
| CDK12 | Uterus–AdenoCa | serous ovarian | TSG |
| CDK4 | CNS–GBM | | oncogene |
| CIITA | Lymph–BNHL | PMBL, Hodgkin lymphoma | TSG, fusion |
| CNTNAP2 | CNS–GBM | glioma, melanoma | TSG |
| CSMD3 | Lymph–CLL | ovarian cancer, oral SCC, lung cancer | TSG |
| CTNNA2 | Lung–SCC | gastric cancer | oncogene |
| CTNND1 | Breast–AdenoCa | large intestine carcinoma | |
| CTNND2 | Head–SCC | prostae adenocarcinoma, GIST | oncogene |
| CXCR4 | Lymph–CLL, Lymph–BNHL | WM | oncogene |
| DDIT3 | CNS–GBM | liposarcoma | oncogene, fusion |
| EGFR | CNS–GBM | glioma, NSCLC | oncogene |
| EPHA3 | Liver–HCC | lung cancer, CRC, melanoma | |
| EPS15 | Biliary–AdenoCa | ALL | TSG, fusion |
| ERBB4 | Eso–AdenoCa | melanoma, gastric, NSCLC | oncogene, TSG |
| FAM135B | Lung–SCC, Head–SCC | SCLC | |
| FHIT | Lymph–BNHL | pleomorphic salivary gland adenoma | TSG, fusion |
| GRM3 | Stomach–AdenoCa, CNS–GBM | melanoma, oral SCC | oncogene |
| HIF1A | Thy–AdenoCa | endometrioid carcinoma, glioblastoma, colorectal, renal, lung, pancreatic | oncogene |
| HOOK3 | Prost–AdenoCa | papillary thyroid | fusion |
| IKZF1 | CNS–GBM | ALL, DLBCL | TSG, fusion |
| IRF4 | Breast–AdenoCa, Prost–AdenoCa, Skin–Melanoma, Uterus–AdenoCa, Eso–AdenoCa, Stomach–AdenoCa, CNS–GBM, Lung–SCC, ColoRect–AdenoCa, Biliary–AdenoCa, Lung–AdenoCA, Liver–HCC | MM | oncogene, TSG, fusion |
| KDR | Liver–HCC | NSCLC, angiosarcoma | oncogene |
| KDSR | Lymph–CLL, Lymph–BNHL | B–NHL | fusion |
| LPP | Lymph–BNHL | lipoma, leukaemia | oncogene, fusion |
| LRIG3 | CNS–GBM | NSCLC | TSG, fusion |
| LRP1B | Eso–AdenoCa | CLL, ovarian cancer, oesophageal squamous cell carcinoma, urothelial cancer | TSG |
| MACC1 | Liver–HCC | liver, CRC | oncogene |
| MECOM | Skin–Melanoma | AML, CML, MDS | oncogene, fusion |
| MLLT6 | Thy–AdenoCa | AL | fusion |
| MYC | Lymph–BNHL | Burkitt lymphoma, amplified in other cancers, B–CLL | oncogene, fusion |
| N4BP2 | Lymph–BNHL | lung cancer | TSG |
| NBEA | Stomach–AdenoCa | large intestine carcinoma, multiple myeloma | |
| NCOA2 | Breast–AdenoCa | AML, chondrosarcoma, rhabdomyosarcoma | oncogene, fusion |
| NRG1 | CNS–GBM | NSCLC | TSG, fusion |
| PAX5 | Lymph–BNHL | NHL, ALL, B–ALL | oncogene, TSG, fusion |
| PDGFRA | Thy–AdenoCaA | GIST, idiopathic hypereosinophilic syndrome, paediatric glioblastoma | oncogene, fusion |
| PIK3CA | Breast–AdenoCa | colorectal, gastric, glioblastoma, breast | oncogene |
| PREX2 | Eso–AdenoCa | melanoma, pancreatic ductal adenocarcinoma | oncogene |
| PTPRC | Stomach–AdenoCa | T–ALL | TSG |
| PTPRT | Skin–Melanoma, Eso–AdenoCa | HNSCC, colorectal cancer, gastric cancer, lung cancer, melanoma | TSG |
| QKI | ColoRect–AdenoCa | angiocentric glioma, colorectal cancer | oncogene, TSG |
| RAD51B | Lymph–BNHL | lipoma, uterine leiomyoma | TSG, fusion |
| RGS7 | Breast–AdenoCa, Lung–SCC, Head–SCC | melanoma | |
| RHOH | Lymph–BNHL | NHL | TSG, fusion |
| RMI2 | Lymph–BNHL | PMBL, Hodgkin lymphoma | TSG, fusion |
| RNF43 | Lymph–BNHL | cholangiocarcinoma, ovary, pancreas | TSG |
| ROBO2 | ColoRect–AdenoCa | colorectal adenocarcinoma, melanoma | TSG |
| S100A7 | Breast–AdenoCa | melanoma, T-cell lymphoma | fusion |
| SFRP4 | Eso–AdenoCa, Stomach–AdenoCa | colorectal cancer, melanoma, SCC, gastric cancer, oesophageal SCC | TSG |
| SGK1 | Lymph–BNHL | Nodular lymphocyte predominant Hodgkin lymphoma | oncogene |
| SOCS1 | Lymph–BNHL | Hodgkin lymphoma, PMBL | TSG |
| TCL1A | Lymph–CLL, Lymph–BNHL | T–CLL | oncogene, fusion |
| TERT | CNS–GBM, Thy–AdenoCa | melanoma, glioblastoma, hepatocellular carcinoma, bladder, skin basal cell, skin squamous cell, mesothelioma, medulloblastoma, other tumour types | oncogene, TSG |
| TOP1 | Thy–AdenoCa | AML* | fusion |
| TRIP11 | Head–SCC | AML | fusion |

**Supplementary Figure 5.** Known cancer genes identified in the 100-kbps genomic regions with mutation burden significantly exceeding random forest predictions informed by CA and RT profiles.

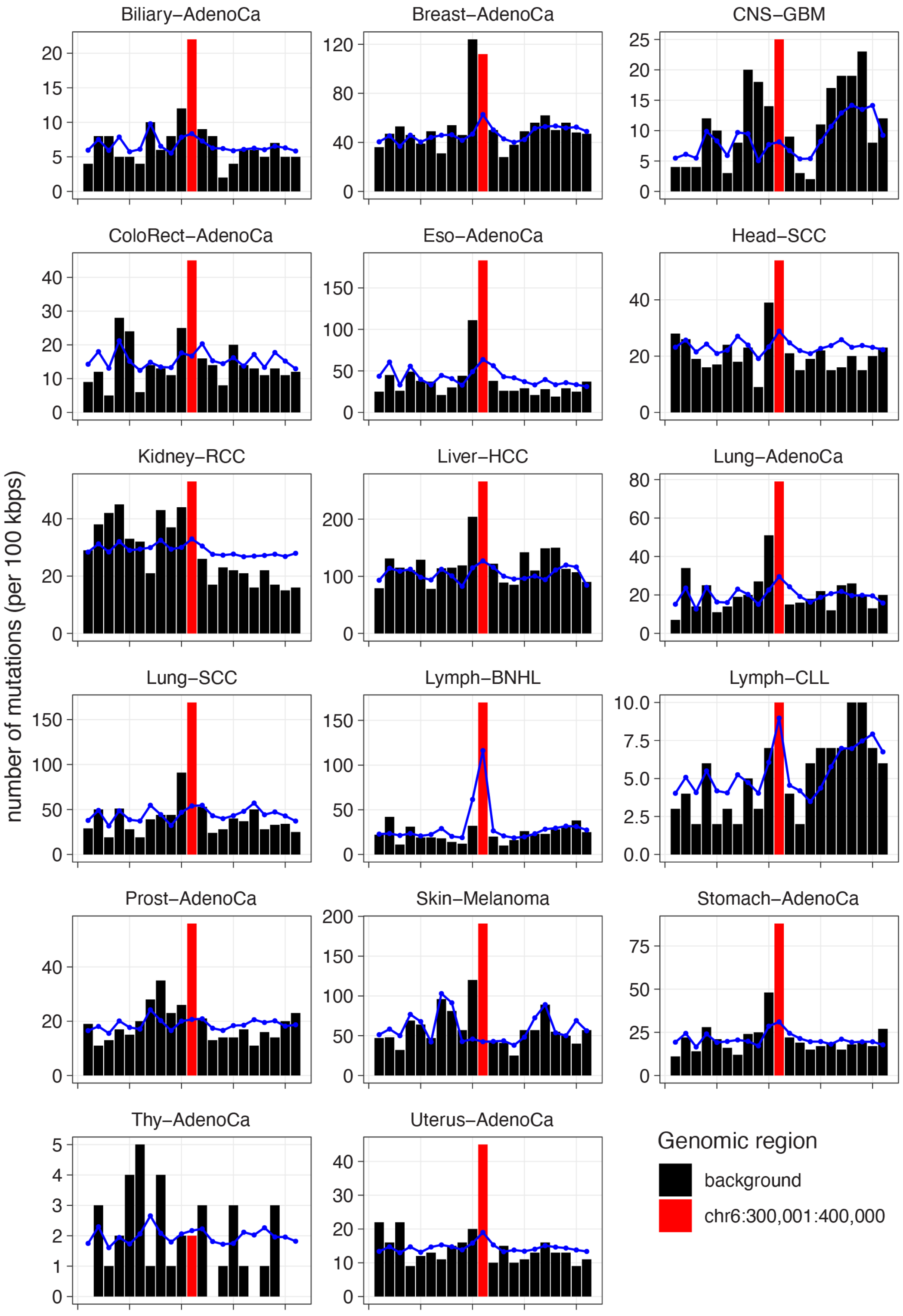

**Supplementary Figure 6.** Regional mutation burden of the frequently mutated region encoding *IRF4* and *DUSP22* (red) and the 20 adjacent regions (black). Bars represent 100-kbps regions. Predicted mutation burden from random forest models is shown in blue.
